# The Parkinson’s disease associated Leucine-rich repeat kinase 2 affects expression of Transferrin receptor 1 and phosphorylation of key signaling proteins in human iPSC-derived dopaminergic neurons

**DOI:** 10.1101/2025.05.06.652393

**Authors:** Natalja Funk, Felix Knab, Marita Munz, Thomas Ott, Thomas Gasser, Saskia Biskup

## Abstract

Several variants in the *Leucine-rich repeat kinase 2* (*LRRK2*) gene account for familiar and sporadic late-onset Parkinson’s disease (PD). LRRK2 is a large, multifunctional kinase involved in different intracellular pathways crucial for homeostasis and cell survival. One of the poorly understood mechanisms of Parkinsonism is the iron accumulation in *Substantia nigra pars compacta* (*SNc*). Transferrin receptor 1 (TfR) plays a significant role for iron uptake into the cell. Here, we investigated the expression of TfR in human induced pluripotent stem cell (iPSC)-derived dopaminergic neurons (hDANs) generated from PD patients carrying the LRRK2 p.G2019S form and found dysregulated TfR levels. In addition, we found gene status depending variations of LRRK2 expression in differentiated hDANs, while neuronal progenitor cells (NPCs) did not display these changes. This suggests an unknown regulatory mechanism of LRRK2 expression during dopaminergic differentiation. Further investigations showed dysregulated phosphorylation of the PD-associated GSK-3β und the key signaling factor Akt.

## Introduction

Parkinson’s disease (PD) is the most prevalent neurodegenerative disorder after Alzheimer’s disease. It affects ∼1-2% of the population over 60 years of age, causing characteristic motor dysfunctions and affecting autonomic functions and cognition. PD is histopathologically characterized by progressive degeneration of midbrain dopaminergic (mDA) neurons in the *Substantia nigra pars compacta* (*SNc*) and the presence of cytoplasmic inclusions (Lewy bodies) (Schapira et al., 2017). Most cases are considered idiopathic, only a minor percentage of PD patients (about 5-10%, dependent on population) are known to have familiar forms (Bandres-Ciga et al., 2020).

One of the most common genetic risk factors in late-onset PD are variants within the *Leucine-rich repeat kinase 2* (*LRRK2*). Several mutations in the *LRRK2* gene account for ∼4% of familial PD up to 36% in some populations and up to 2% of sporadic late-onset Parkinsonism (Healy et al., 2008, Li et al., 2014; Lin et al., 2014, Ferreira et al., 2016, Sosero et al., 2023). LRRK2 is a multi-domain protein involved in a variety of cellular processes crucial for homeostasis and cell survival in neuronal and non-neuronal cells. Multiple LRRK2-mediated mechanisms have been proposed in PD pathophysiology including apoptosis, mitochondrial and lysosomal function, autophagy, synaptic transduction/vesicle formation, inflammation, and several others (Jeong et al., 2020, Sosero et al., 2023). Furthermore, a number of proteins including Rab family GTPases, EndophilinA, Rac1, 14-3-3 proteins and many others have been described as putative substrates and interactors of LRRK2 *in vivo* and/or *in vitro* (Steger et al., 2016, Seol et al., 2019, Jeong et al., 2020).

The potential role of iron metabolism in the pathophysiology of Parkinsonism has a long history. Markedly increased levels of iron in the *SNc* of PD patients were initially described in the 1920’s (Lhermitte et al., 1924, Foley et al., 2022). To date, almost 100 years later, the mechanisms responsible for this phenomenon are still poorly understood. Also, the question, whether a change in iron metabolism is a cause or consequence of PD, remains unsolved. Controlling iron levels at systemic and cellular levels is a crucial part of many aspects of human health and diseases (Mochizuki et al., 2020). A potential role of iron in PD pathogenesis is supported by the fact that mutations in many genes related to iron homeostasis are associated with a higher relative PD-risk (reviewed in Rhodes and Ritz, 2008).

The iron homeostasis in human comprises systemic and cellular iron regulation. The regulation at the cellular level includes iron import, intracellular iron pools (labile and storage), iron export and translational control (summarized in Campos-Escamilla, 2021, Ma et al., 2021). Most cells take up iron through receptor-mediated endocytosis via Transferrin receptor 1 (TfR, also known as Cluster of Differentiation 71 (CD71)), a transmembrane glycoprotein composed of two disulfide-linked monomers (summarized in Aisen, 2004). In the CNS, the TfR is described to be mainly located in endothelial cells. Moreover, TfR immunoreactivity was detected intraneuronally in several brain regions without access to peripheral blood: the TfR expressed cells were mainly confined to the cerebral cortex, hippocampus, habenular nucleus, red nucleus, pontine nuclei, reticular formation, several cranial nerve nuclei, deep cerebellar nuclei, the cerebellar cortex and the *Substantia nigra* (Moos, 1996). Astrocytes, oligodendrocytes, or microglial cells do not show TfR immunoreactivity (Moos, 1996). The molecular mechanism of the TfR-mediated endocytosis has already been well investigated (Campos-Escamilla, 2021). Several members of the Rab GTPase family have been shown to regulate distinct steps in the intracellular TfR cycle. As described by Matsui et al. (2011), small GTPase Rab12 is mainly localized at TfR-positive recycling endosomes and is specifically involved in degradation of TfR and consequently regulated its level in the cell. The function of Rab12 is controlled by GDP-GTP exchange factor DENND3 (Yoshimura et al., 2010) and potentially through phosphorylation by the PD-linked LRRK2 (Steger et al., 2016, Thirstrup et al., 2017).

In this work, we investigated the possible link between the pathogenic *LRRK2* c.6055G>A (G2019S) variant and TfR expression in human induced pluripotent stem cell (iPSC)-derived dopaminergic neurons (hDANs) generated from two PD patients. We found that the *LRRK2* c.6055G>A (G2019S) form is associated with dysregulation of LRRK2 and TfR protein expression levels in hDANs. To exclude a cell-model dependent unspecific effect observations, we further studied the effect of altered LRRK2 expression on TfR protein level in fibroblasts from Lrrk2-deficient and transgenic human LRRK2 p.G2019S-overexpressed rats. Consistent with our previous findings, we found that the Lrrk2 expression level affects the absolute level of TfR protein expression in this cell type. We further found that the phosphorylation of serine-threonine kinase glycogen synthase kinase-3 beta (GSK-3β) and the key signaling factor protein kinase B (PKB, also known as Akt) was significantly changed in *LRRK2* c.6055G>A (G2019S) mutated hDANs.

Interestingly, our notably observation was that the pathogenic *LRRK2* c.6055G>A variant do not affect the LRRK2 protein expression level in iPSC-derived neural progenitor cells (NPCs) of PD patients. Furthermore, we found completely different regulation of protein expression and phosphorylation in examined NPCs. Collectively, our data demonstrates a role of the Parkinson’s disease-linked LRRK2 in TfR expression and regulation of diverse regulatory proteins.

## Results

### LRRK2- and TfR-expression levels are affected in G2019S mutated hDANs

To evaluate whether LRRK2 p.G2019S mutation could influence the expression of TfR in neuronal cells, we cultured iPSC-derived hDANs generated from one early-onset LRRK2-associated PD patient (referred to as L1-1Mut) and one late-onset LRRK2 carrier with PD (L2-2Mut) and their corresponding isogenic gene-corrected cell lines (designated L1-1GC and L2-2GC, control lines where the mutation *LRRK2* c.6055G>A (G2019S) was corrected to the consensus sequence). 23 days after induced dopaminergic differentiation, the neuronal maturity of the cells was analyzed by immunocytochemistry (ICC) using microtubule associated protein 2 (MAP2) and tyrosine hydroxylase (TH) as markers. Roughly 75% of the differentiated L1-1 cells and 91% of L2-2 cells were positive for MAP2 while 30-40% of the cells expressed TH (a detailed evaluation of this ICC data as well as further analysis of the cells is described in Knab et al., 2025).

Next, the expression of TfR in iPSC-derived hDANs was analyzed by ICC and Western blotting. Using immunofluorescence, we indicated that the cells expressing dopaminergic marker TH and neuron-specific marker β-tubulin class III showed characteristic TfR puncta-like staining (Fig. 1a). By Western blot we found significant differences in TfR protein levels between G2019S-mutated and gene corrected hDANs: the expression was upregulated in L1-1Mut cells in comparison to L1-1GC control line (2.9-fold, Fig. 1b, p = 0.005) and downregulated in L2-2Mut line compared to its gene corrected L2-2GC variant (3.4-fold, Fig. 1c, p = 0203, Mann-Whitney U-test). Interestingly, the parallel analysis of LRRK2 expression in hDANs also showed significant differences between mutated and gene corrected cells: the expression level was upregulated in L1-1Mut (2.5-fold, Fig. 1b) and strongly downregulated in L2-2Mut line (Fig. 1c).

**Fig. 1.**
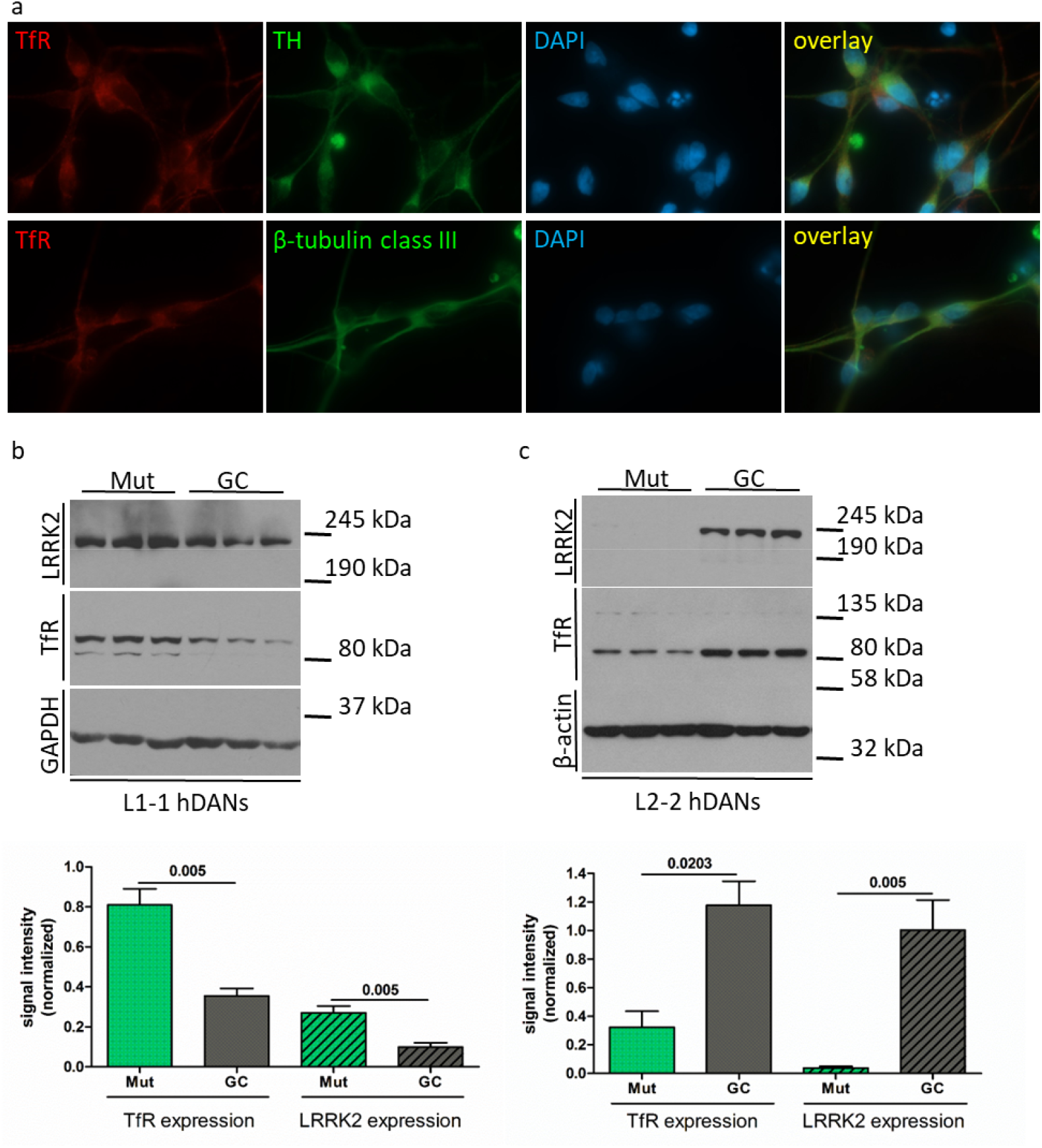
Expression of TfR in human iPSC-derived cells. (a) Human iPSC-derived TH-immunoreactive cells (top) and β-tubulin class III positive neurons (button) express TfR. (b and c) Western blot analysis (top) and quantification (button) of LRRK2- and TfR-protein expression in L1-1 hDANs (b) and L2-2 hDANs (c), n = 3 independent differentiation experiments.

### Expression of Lrrk2 affects the TfR expression level in rat fibroblasts

Based on our data from hDANs, we asked whether the dysregulation of the TfR expression was related to the changes in LRRK2 protein level. Therefore, we investigated the TfR expression in fibroblasts from Lrrk2-deficient (referred to as Lrrk2 k.o.) and human LRRK2 p.G2019S-transgenic (named as hG2019S tg) rats generated in our lab (Suppl. Fig. 1). As described in Funk et al. (2019), Lrrk2 k.o. animals lacking Lrrk2 protein expression in all tissues tested, including brain, spleen, and kidney. hG2019S tg rats express human LRRK2 p.G2019S in different tissues (Suppl. Fig. 1) and cell types (including fibroblasts), as published in Walker et al. (2014). As shown by Western blot analysis, the expression of TfR is significantly reduced in Lrrk2-deficient rat fibroblasts (p-value 0.005) in comparison to wild-type cells (Fig. 2a). The overexpression of human LRRK2 p.G2019S results in a slight, but not significant upregulation of TfR level in hG2019S tg rat fibroblasts (p = 0.582, Fig. 2a).

**Fig. 2.**
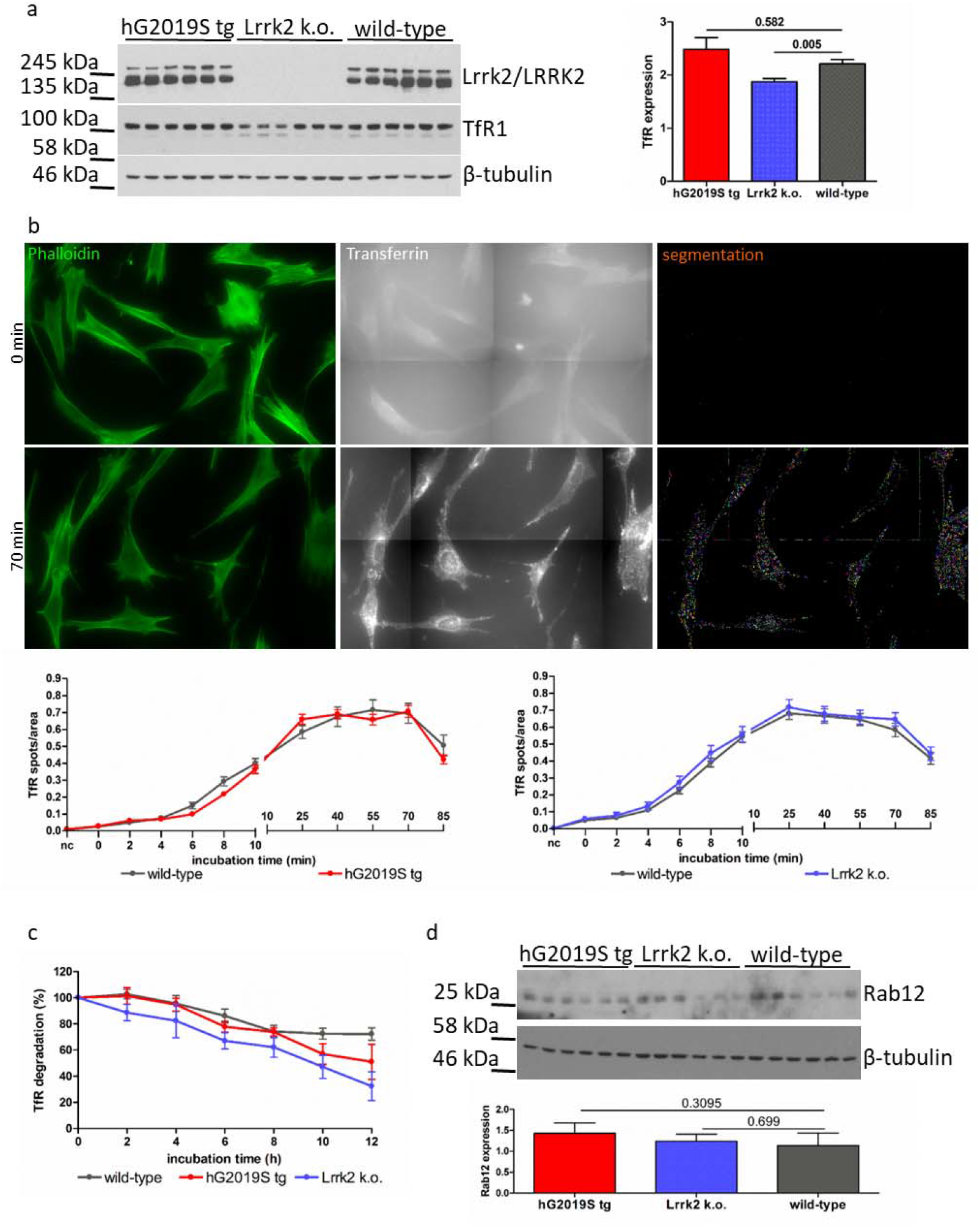
TfR in rat fibroblasts. (a) Western blot analysis and quantification of TfR protein expression in hG2019S-transgenic (hG2019S tg), Lrrk2-deficient (Lrrk2 k.o.) and wild-type fibroblasts (n = 8 independent experiments). (b) Transferrin recycling assay in rat fibroblasts: Phalloidin (green), DAPI (data not shown) and TexasRed transferrin (white) staining were used for automated quantification of generated data (segmentation and ROI count). Representative images for wild-type fibroblasts, time points 0 und 70 min (top). Graphical representation of transferrin uptake at different time points (= incubation time) in hG2019S tg (left, n = 11 independent experiments) and Lrrk2 k.o. (right, n = 9 independent experiments) fibroblasts (button). (c) TfR protein degradation at different time points after cycloheximide addition (n = 8 independent experiments). (d) Western blot analysis/ quantification of Rab12 expression in hG2019S tg, Lrrk2 k.o. and wild-type fibroblasts.

Most cells, including fibroblasts and neurons, take up iron through receptor-mediated endocytosis via TfR. To investigate the impact of TfR downregulation on its internalization, we observed the TfR translocation by Transferrin Recycling Assay using the TexasRed-labeled transferrin and BD Pathway 855 imaging system in the same rat cell lines. After the addition of transferrin, the internalization of transferrin-TfR complexes occurred and they accumulated in the cytoplasm (Fig. 2b). Despite alterations in total TfR expression, we did not find any differences in transferrin uptake between Lrrk2 k.o., human LRRK2 p.G2019S-transgenic and wild-type fibroblasts (Fig. 2b).

The reduced TfR-level in Lrrk2-deficient rat fibroblasts and human iPSC-derived dopaminergic neurons L2-2Mut can be caused by its lower production or/and increased degradation. As described by Matsui et al. (2011), constitutive degradation of TfR is regulated by small GTPase Rab12, a putative phosphorylation substrate of LRRK2 (Steger et al., 2016, Thirstrup et al., 2017). To investigate the dependency of the TfR stability on LRRK2 expression, rat fibroblasts were treated with cycloheximide, an inhibitor of protein synthesis. This treatment of the cells caused a reduction in the amount of TfR protein in a time-dependent manner, with the estimated half-life over 12 hours in rat fibroblasts. Using the Western blot analysis, we did not find significant differences in degradation of TfR between Lrrk2-deficient, human LRRK2 p.G2019S-transgenic and wild-type cells (Fig. 2c). The quantification of Rab12 level showed similar expression in all investigated rat cell lines (Fig. 2d), the analysis of endogenous Rab12-phosphorylation by SDS-PAGE and Phos-tag electrophoresis followed by Western blot failed in our hands due to weak Rab12 expression in fibroblasts (data not shown).

### The phosphorylation of GSK-3β and Akt is altered in G2019S mutated hDANs

Several proteins have been identified as putative substrates for LRRK2 (Steger et al., 2016, Seol et al., 2019, Jeong et al., 2020). Furthermore, LRRK2 is described to be involved in a variety of cellular pathways (Jeong et al., 2020, Sosero et al., 2023). Therefore, based on our data for altered LRRK2 expression level in hDANs, we analyzed the phosphorylation of the Parkinson’s disease-associated GSK-3β and the key signaling protein Akt in L1-1 and L2-2 lines. Consistent with our LRRK2 expression findings, we found significant alterations in the phosphorylation of GSK-3β and Akt in hDANs (Fig. 3). The phosphorylation of both proteins was upregulated in mutated L1-1Mut cells (P-GSK-3β S9 p = 0.0043, P-Akt T308 p = 0.0022, P-Akt S473 p = 0.0152) and downregulated in L2-2Mut cells (P-GSK-3β p = 0.002, P-Akt S473 p = 0.0022), compared to its gene corrected lines L1-1GC and L2-2GC respectively. The total expression of GSK-3β as well as Akt was not dependent on LRRK2 expression level (two-tailed Mann-Whitney test, Fig. 3). We therefore concluded that the regulation (but not the expression) of several multifunctional signaling factors is altered in LRRK2 p.G2019S hDANs.

**Figure 3.**
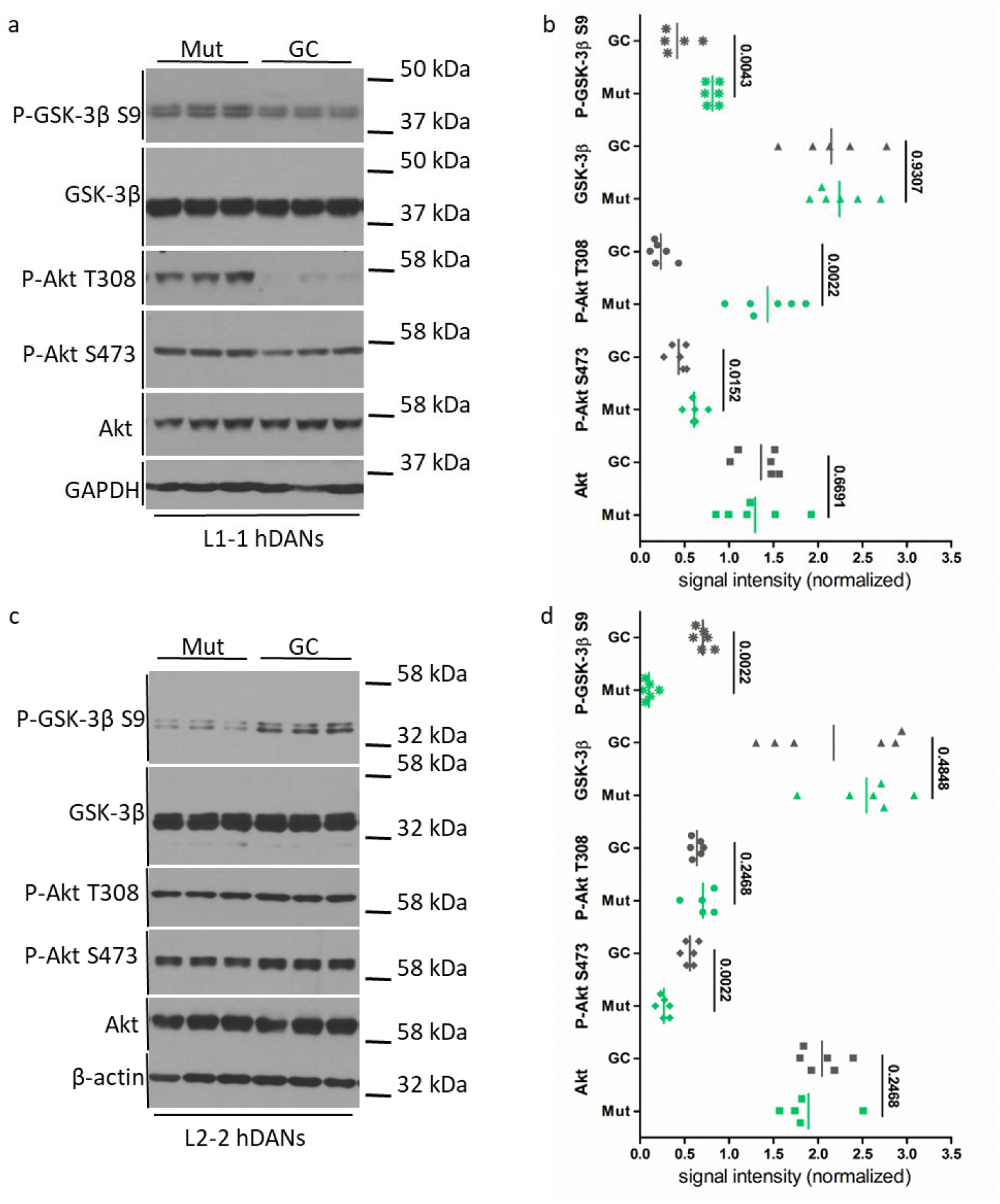
Phosphorylation of GSK-3β and Akt in hDANs. (a) Western blot analysis and (b) quantification of GSK-3β and Akt protein expression and phosphorylation in L1-1 hDANs (n = 3 independent differentiation experiments). (c) Western blot analysis and (d) quantification of GSK-3β and Akt protein expression and phosphorylation in L2-2 hDANs (n = 3 independent differentiation experiments).

### NPCs show no differences in the LRRK2 expression level

hDANs used in this study were generated by induced dopaminergic differentiation of iPSC-derived neural progenitor cells. As previously shown, the LRRK2 expression level was altered in hDANs carrying *LRRK2* c.6055G>A (G2019S) mutation. We asked whether this *LRRK2* variant influence its own expression already in progenitor cells before the dopaminergic differentiation start. Using Western blotting method, we analysed protein extracts from L1-1 and L2-2 NPC lines and found comparable levels of LRRK2 expression in both G2019S-mutated and gene corrected NPCs (L1-1 p = 0.8182, L2-2 p = 0.1255, Fig. 4). Furthermore, following our previous observations in hDANs, we investigated the expression of TfR protein and the phosphorylation of GSK-3β and Akt. Surprisingly, we found a completely different TfR regulation behavior in cell lines studied: both L2-2 NPC lines (L2-2Mut and L2-2GC) showed a very similar expression of TfR (p = 0.3095); the L1-1Mut line expressed reduced level of TfR (p = 0.0022) compared to L1-1GC cells (Fig. 4). The phosphorylation on Ser9 of GSK-3β was not significantly different between mutated and gene corrected L2-2 NPCs (p = 0.8182), and upregulated in early-onset L1 patient (p = 0.0051, two-tailed Mann-Whitney test, n = 3 independent experiments, Fig. 4). The cells from both patients showed strong upregulation of Akt on the threonine 308 (L1-1 p = 0.0079, L2-2 p = 0.0022) with comparable expression of total Akt (Suppl. Fig. 2).

**Figure 4.**
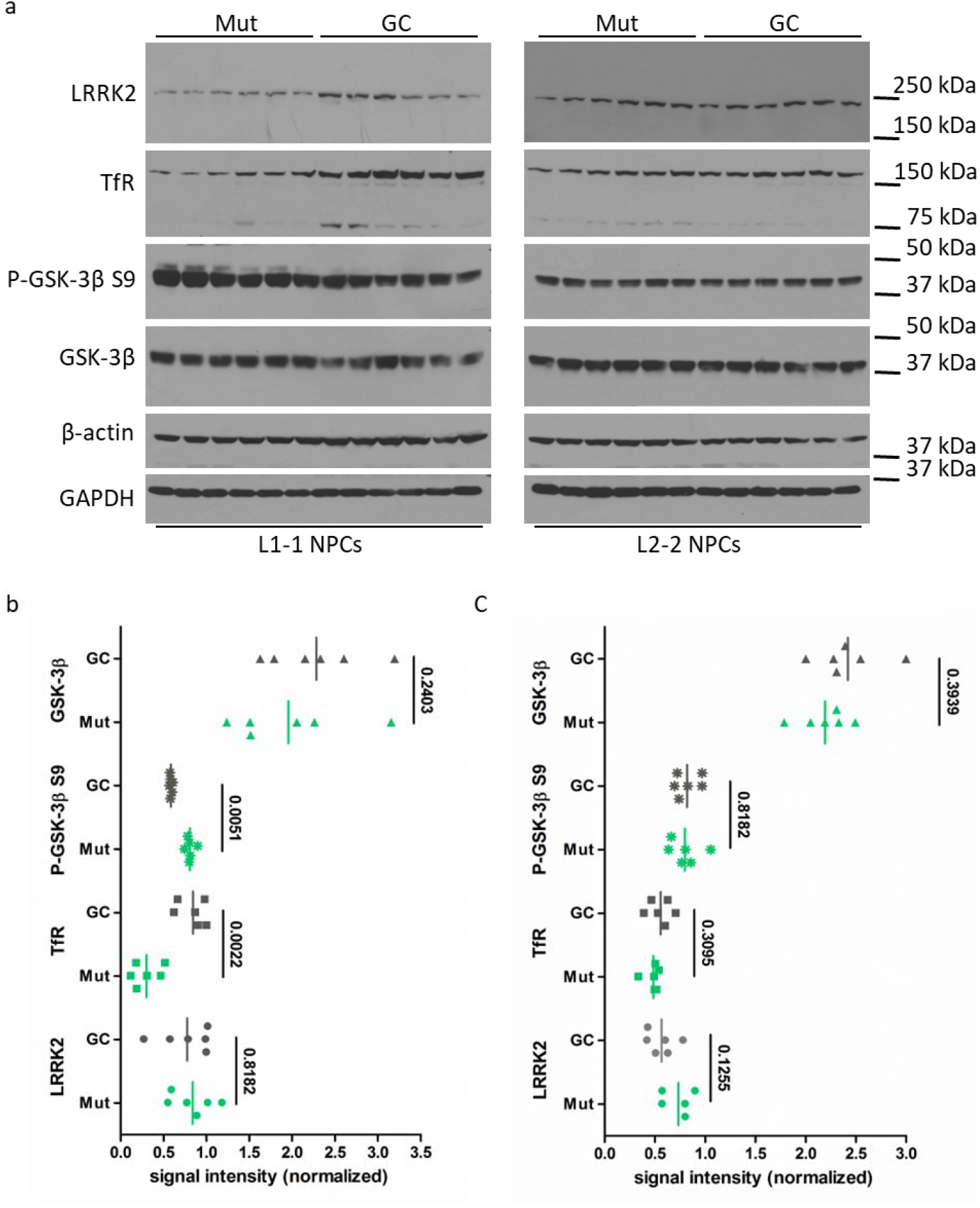
Protein expression in human NPCs. (a) Western blot analysis of LRRK2 and TfR protein expression and GSK-3β phosphorylation in L1-1 NPCs (left) and L2-2 NPCs (right). (b) Quantification of previous Western blot data for L1-1 and (C) L2-2 NPCs (n = 3 independent experiments).

## Discussion

Iron is an abundant element in all organisms, possessing an essential physiological role due to its simplicity in donating and accepting electrons (interconversion between Fe^2+^ and Fe^3+^ forms). The iron distribution in a healthy brain appears to be quite heterogeneous; in brains of patients with neurodegenerative diseases the accumulation of iron in specific affected brain regions have been observed. Various studies reported increased iron deposition in melanin containing DA neurons of the *SNc* of PD patients, the main site of neuronal degeneration (Dexter et al., 1987, Riederer et al., 1989, Götz et al, 2004, Gerlach et al., 2006). However, the specific role of iron in the pathobiology of Parkinsonism is complex and poorly understood. In particular, the exact molecular damage within the *SNc* of PD brain that leads to iron accumulation also remains unclear.

Within this study we investigated the possible role of the Parkinson’s disease-linked LRRK2 in cellular iron homeostasis. Using human iPSC-derived dopaminergic neurons from PD patients with the known pathogenic *LRRK2* c.6055G>A (G2019S) mutation and corresponding isogenic control lines we showed that LRRK2 plays a crucial role in expression of Transferrin receptor 1, a transmembrane glycoprotein important for the iron uptake in neuronal and non-neuronal cells. Furthermore, using the standard Western blot analysis we found a strong dysregulation of LRRK2 expression in G2019S-mutated hDANs: the LRRK2 protein level is strongly elevated in L1-1Mut and reduced in L2-2Mut cells. The expression of TfR seems to be dependent on total LRRK2 amount in a cell: the up- or down-regulation of LRRK2 is accompanied by dysregulation of the TfR expression in the same direction. Additional investigations using Lrrk2-deficient und human LRRK2 p.G2019S-transgenic rat fibroblasts confirmed this observation: the expression of TfR was reduced in Lrrk2 k.o. cells, the overexpression of human LRRK2 p.G2019S lead to a slight but non-significant upregulation of TfR level. Although the effect of LRRK2 dysregulation on TfR expression in rat fibroblasts is not as pronounced as in hDANs, all this data taken together support the assumption of a connection.

Changes in expression of a protein raise a number of possible questions. Therefore, we observed the TfR translocation using the fluorescence-labeled transferrin and BD Pathway 855 imaging system. Because of their flat morphology, fibroblasts are a very good candidate for this type of study; in contrast, hDANs have small cell bodies, form aggregates and are poorly suitable for this. On the other hand, the expression of TfR in investigated hDANs is clearly more impaired than in fibroblasts. Due to the biological and technical limitations we used rat fibroblasts for the transferrin uptake study and did not find any effect of TfR dysregulation in Lrrk2-deficient and hG2019S tg cells. Matsui et al. (2011) described the Rab12 GTPase, a putative substrate of LRRK2 (Steger et al., 2016, Thirstrup et al., 2017), as a crucial regulator of constitutive degradation of TfR. It is conceivable that the alterations in the TfR expression are due to changes in the regulation of Rab12 by LRRK2. Therefore, we analysed the TfR stability in Lrrk2 k.o. and hG2019S tg rat fibroblasts by treatment with cycloheximide over 12 hours and did not find significant differences in TfR degradation in this cell type. However, because fibroblasts in general are not an ideal cell model to study the physiology of dopaminergic neurons, the interpretation of our findings around the internalization and degradation of TfR is strong limited.

Ohta et al. (2015) described lower LRRK2 protein level in induced iPSC-derived neurons from fibroblasts of PD patients with LRRK2 p.I2020T in the Sagamihara family. This observation is very similar to our findings in L2-2Mut hDANs with LRRK2 p.G2019S variant. Furthermore, the authors found a dysregulation of Akt phosphorylation and a reduction of phosphorylation at Ser9 of GSK-3β / an increased phosphorylation at Tyr219 of GSK-3β in LRRK2 p.I2020T mutated cells (Otha et al., 2015). These published data are consistent with our results for the L2-2 hDANs too: using Western blot analysis we found strong downregulation of the inhibited GSK-3β phosphorylation at Ser9 and reduced phospho-Ser473 Akt signal. In the L1-1Mut hDANs, the results are almost mirror-inverted: the LRRK2 protein expression and the phosphorylation of Akt and GSK-3β Ser9 were upregulated. GSK-3β has been identified to be involved in several intracellular signalling pathways, including but not limited to Wnt/β-catenin, brain-derived neurotrophic factor (BDNF), insulin and Hedgehog signalling, and the translation initiation factor eIF-2B (summarized in Wildburger and Laezza, 2012). In particular, the crucial role of GSK-3β in the phosphorylation of Tau makes this protein attractive to neurobiologists. GSK-3β is constitutively active in cells (Doble and Woodgett, 2003) and negatively regulated by Akt through phosphorylation of Ser9 (Cross et al., 1995, Strivastava and Pandey, 1998). In addition, LRRK2 may function as an enhancer for GSK-3β (Kawakami et al, 2014). Taken together we speculated that LRRK2 expression affect the phosphorylation of Akt and, as a result, the activity of GSK-3β. However, the question about the origin for the dysregulated LRRK2 protein expression remains unclear. Published studies demonstrated impaired stability for the pathogenic LRRK2 p.I2020T variant and LRRK2 p.A1442P and R1441C ROC domain mutants (Otha et al., 2009, Greene et al., 2014). In contrast, the degradation rate of LRRK2 p.G2019S mutant was not different from the wild-type form (Otha et al., 2009). Increasing evidence suggests that neurodevelopmental alterations might increase the risk to become neurodegenerative diseases. Therefore, we analysed the LRRK2 expression in iPSC-derived neural progenitor cells, the cells that were used to generate hDANs for this study. Our data indicate that the LRRK2 expression in iPSC-derived NPCs (= before the dopaminergic differentiation starts) is not dependent on genetic *LRRK2* constellation: the LRRK2 p.G2019S mutated and corresponding gene corrected isogenic hDANs expressed comparable amount of LRRK2 in both L1-1 and L2-2 lines.

The modifications of the spatial chromatin organization and dynamic epigenetic landscapes of cells are discussed as important regulators during differentiation. Epigenetic modifications in enhancers, often looped with target promotors, can reprogram the transcription of cell-type specific genes (Heintzman et al., 2009, Lin et al., 2022). Meléndez-Ramírez et al., (2021) published transcriptomic changes and chromatin accessibility during differentiation of embryonic stem cells into dopaminergic neurons. However, it remains unknown how the single nucleotide polymorphisms (SNPs) influence the 3D genome. Lin et al., (2022), demonstrated that SNP rs1873613 upregulate the expression of the *LRRK2* gene during immune activation of macrophages. This SNP is located in an enhancer upstream of *LRRK2* and interacts with *LRRK2* promoter through a long-range chromatin loop (Lin et al., 2022). Oliveira et al., (2021), described *LRRK2* variants associated with alcohol dependence in multiethnic populations from South and North America. Interestingly, these SNPs are located in non-coding regions of LRRK2, partly in Regulatory Build (Oliveira et al., 2021). In this study we found a dysregulation of LRRK2 level in LRRK2 p.G2019S expressing hDANs in comparison to isogenic control cells. The LRRK2 expression in progenitors of these neurons is indistinguishable between mutated and control lines. Taken together, we speculate that variants within coding regions of *LRRK2* can modulate itself expression during dopaminergic differentiation, and the interaction of this variants with genetic background would affect the regulation of *LRRK2* gene in an unknow way, possibly by reorganization of the spatial chromatin conformation.

Our additional investigations in NPCs demonstrated a strong elevation of phosphorylation of Akt on T308 in both LRRK2 p.G2019S lines. This upregulation seems to be not dependent on total amount of LRRK2, because all mutated and control cells expressed comparable level of LRRK2. The phosphorylation of GSK-3β on Ser9 is described to be negatively regulated by Akt (Cross et al., 1995, Strivastava and Pandey, 1998). In NPCs, the phosphorylation of GSK-3β was upregulated in L1-1 Mut line and not affected in L2-2 line. The expression of TfR is differently regulated in analysed cells too: it was increased in L1-1 Mut and not affected in L2-2 Mut lines in comparison to corresponding isogenic controls. Naively, we can assume that the cells from the LRRK2-associated PD patient L1 contains additionally regulatory modifiers, which can lead to an earlier onset of the disease (age of onset is 40.36, Suppl. Table 1). Genome-wide association studies (GWAS) have revealed thatL>L80% of disease-associated SNPs are found within poorly annotated regions, making the identification of causal polymorphisms a difficult task (Buniello et al., 2019, Meléndez-Ramírez et al., 2021). Further studies are needed to explore the exact mechanisms behind these observations.

Collectively, we conclude that LRRK2 contributes to regulation of TfR expression in human iPSC-derived dopaminergic neurons. Furthermore, we assume that *LRRK2* variants influence expression of its own during dopaminergic differentiation by an unknown mechanism. In addition, based on our data from iPSC-derived cells and corresponding PD patients, we speculated that dependent on the genetic background the expression of LRRK2 during dopaminergic differentiation can be up- or down-regulated, which co-determined the disease onset.

## Experimental Procedures

### Ethics statement and data protection

All procedures were in accordance and approved by the ethical board at the University of Tuebingen and according to the international standards defined in the declaration of Helsinki. Human samples were obtained with consent and prior ethical approval at The University of Tuebingen and the Hertie Institute for Clinical Brain Research Biobank number 146/2009B01 and the Medical Faculty of the University of Tübingen (https://www.medizin.uni-tuebingen.de/de/medizinische-fakultaet/ethikkommission).

### Derivation of human neural progenitor cell (NPC) lines and differentiation into dopaminergic neurons (hDANs)

iPSCs from dermal fibroblasts of two PD patients harbouring the *LRRK2* c.6055G>A mutation (referred to as L1 and L2, Table S1) as well as gene-corrected iPSC lines were generated as described in Reinhardt et al., 2013a. Individual clonal iPSC lines derived from both patients were designated L1-1Mut and L2-2Mut (carrying *LRRK2* c.6055G>A (G2019S) mutation) and L1-1GC and L2-2GC (isogenic gene-corrected lines), respectively. The derivation and characterization of iPSC-derived neural progenitor cell lines is reported in Reinhardt et al., 2013b. NPCs were cultured in a “base medium” (containing 50% Neurobasal® medium (Thermo Fisher Scientific, #21103-049), 50% DMEM/F12 (Thermo Fisher Scientific, #11-330-057), 1X Penicillin/Streptomycin (Merck, #A2213), 1X GlutaMax (Thermo Fisher Scientific, #35050-038), 1X B27 supplement (w/o vitamin A; Thermo Fisher Scientific, #12587-010) and 0.5X N2 supplement (Thermo Fisher Scientific, #17502-048)), supplemented with 0.5 µM PMA (Merck, #540220), 3 µM CHIR (Axon Medchem, #Axon1386) and 200 µM ascorbid acid (Sigma-Aldrich, #A4544) on Matrigel (Corning)-coated 6-well cell culture plates.

Differentiation of human NPCs into dopaminergic neurons was done according to published protocols (Bus et al., 2020). Confluent NPCs were incubated in a patterning medium (“base medium” plus the addition of 20 ng/ml of BDNF (PeproTech, #450-02), 10 ng/ml of FGF8 (PeproTech, #100-25), 1 µM of PMA and 200 µM of ascorbic acid) for seven days (days 1 to 7). Then, the cells were matured in maturation medium (“base medium” supplemented with 10 ng/ml of BDNF, 10 ng/ml of GDNF (PeproTech, #450-10), 1 ng/ml of TGF-β3 (PeproTech, #AF-100-36E), 200 µM ascorbic acid, 500 µM dbcAMP (PanReac AppliChem, #A0455) and 10 µM DAPT (Selleckchem, #S2215)) for days 8 to 23.

### Animal models

The Lrrk2 knock-out rats (referred to as Lrrk2 k.o.) were generated in our lab via microinjection of a pair of TAL-nucleases as described in Funk et al., 2019. The rat line used here contains a 7 bp deletion in exon 2 of the *Lrrk2* gene (Suppl. Fig. 1) resulting in an open read frame shift and translational stop (Funk et al., 2019). The hLRRK2 G2019S transgenic rats (referred to as hG2019S tg) were generated in our lab via microinjection of a linearized Bacterial Artificial Chromosome (BAC) containing human genomic G2019S-mutated *LRRK2* region (including regulatory elements) as described in Walker et al., 2014. The BAC clone pBACe3.6 hLRRK2 G2019S was generated in Matthew J. Farrer lab, Vancouver (Walker et al., 2014).

All rat models were kept as heterozygous lines by breeding to CD (SD) rats (Charles River). For Lrrk2 knock-out rats, two independent lines (lines 1 and 2) were kept separate. For experiments, the heterozygous animals from lines 1 and 2 were bred to generate homozygous Lrrk2 deficient rats. hLRRK2 G2019S transgenic rats are used as heterozygous for the transgene. Wild-type littermates were used as wild-type control in all experiments.

### Rat fibroblasts cell culture

Rat fibroblasts were prepared from 6-month-old animals and cultured as described in Funk et al., 2019. To study the TfR stability, the cells were plated on uncoated 24-well culture plates (Nunc, 50 000/ well), cultured overnight, and treated with 50µg/ml cycloheximide (SigmaAldrich/Merck, #01810) for 0-12h before Western blot analysis.

### Transferrin Recycling Assay

To study the TfR-mediated endocytose fibroblasts were seeded on Poly-DL-ornithine hydrobromide (PORN)-coated 96-well microplates (BD Falcon, #353219, 4 000/well) overnight. PORN solution: 0.15 M Boric acid (Merck, #B6768) pH 8.3, containing 0.5 mg/ml Poly-DL-ornithine hydrobromide (SigmaAldrich/Merck, #P0421). After washing with ice-cold RPMI 1640 medium (Biochrom, F1275) the cells were incubated with 0.1 mg/ml Transferrin Texas Red® conjugate (molecular probes, T2875) in RPMI 1640 medium for 30 min on ice, washed again and transferred to 37°C. Afterwards the cells were fixed with paraformaldehyde (Electron Microscopy Sciences, 15710, 4% final concentration) at the times indicated, and washed twice with phosphate buffered saline (PBS Dulbecco, Biochrom, L1825). After permeabilization with 0.1% Triton X100 (Carl Roth, #3051.3) in PBS the cells were incubated with Alexa Fluor® 488 phalloidin (life technologies, A12379, 1U/Well) and DAPI (ThermoFisher Scientific, #62248, 0.5 ng/ml) and washed twice with PBS again. Finally, immunofluorescence was imaged using BD Pathway 855 (settings: Alexa488, DAPI and TexasRed dyes; montage capture setup 5×5 (w/o gap); laser auto-focus). The quantification occurred on AttoVision 1.5 software (BD Bioscience) using two-steps segmentation. Step 1 segmentation setup (=area definition): Segmentation Cyto-NucDual; Shape Dual channel (exclude center); Channel A: DAPI, Channel B: Alexa 488; Threshold manual; Split erosion factor 20; Scrap Object pixels min 300, max -1. Step 2 segmentation setup (ROIs definition): Segmentation Transferrin; Shape polygon; Threshold automatically; Split Watershed, Interations 10; Scrap Object pixels min -1, max 30; Output ROIs Probe 1.

### Western blot analysis

Protein extracts and Western blots were prepared as described in Funk et al., 2019.

### Immunocytochemistry

Mature hDANs were cultured on Matrigel-coated glass coverslips and fixed with 3.7% (w/v) paraformaldehyde (Electron Microscopy Sciences, 15710). For TfR expression analysis, cells were co-stained with anti-TfR and anti-TH or anti-β-Tubulin, class III antibody according to the manufacturer’s protocol for bioimaging. In brief, fixed cells were washed three times with PBS (Biochrom, L1825), permeabilized with 0.1% Triton X-100 (Carl Roth, #3051.3) and blocked with 15% (v/v) NGS in PBS for 1h. For TfR- and TH-staining, cells were incubated with primary antibodies over night at 4°C. After washing with PBS and staining with secondary antibody and DAPI (ThermoFisher Scientific, #62248, 0.5 ng/ml) for 1h at RT, coverslips were mounted using Vectashield mounting medium (Vector Laboratories, H-1500). For TfR- and β-tubulin class III-staining, cells were first stained with anti-TfR antibody over night as described above and finally co-stained with anti-β-Tubulin, class III antibody (1:50) plus DAPI (0.5 ng/ml) and secondary antibody for anti-TfR for 1h at RT.

### Antibodies

Primary antibodies used: Akt #9272 (1:1000), phospho-Akt (Ser473) # 4058 (1:1000), phospho-Akt (Thr308) #2965 (1:1000), GAPDH (D16H11) XP™ #5174 (1:1000), phospho-GSK-3β (Ser9) #9323 (1:1000), GSK-3β (27C10) #9315 (1:1000), β-Tubulin #2146 (1:10000) (all from Cell Signaling Technology); beta-actin [AC-15] ab6276 (1:10000), LRRK2 [MJFF2 (c41-2)] ab133474 (1:500) (abcam); LRRK2 (MJFF5) (1:500) (Michael J. Fox Foundation, Epitomics); Tyrosine Hydroxylase (TH) #AB152 (1:500) (Millipore); Rab12 (H-11) sc-515613 (1:500) (Santa Cruz Biotechnology); Alexa Fluor 488 Mouse anti-β-Tubulin, Class III Clone TUJ1 #560338 (1:50) (BD Pharmingen); Transferrin Receptor (H68.4) #13-6800 (1:2000) (ThermoFisher Scientific).

Secondary antibodies used: anti-rabbit IgG, HRP-linked #7074 (1:5000, Cell Signaling Technology); Goat anti-mouse IgG, HRP-linked #172-1011 (1:10 000, Bio-Rad); Cy™3-conjugated Goat anti-rabbit IgG (1:350, 111-165-046), Cy™2-conjugated Goat anti-mouse IgG (1:350, 115-225-164), Cy™3-conjugated Donkey anti-mouse IgG (1:350, 715-165-150) (all from Jackson ImmunoResearch Laboratories).

### Statistical analysis

All data sets were tested for normal Gaussian distribution using GraphPad Prism 4.02. The data were analysed by two-tailed Mann-Whitney U-test and two-tailed t-test.

## Acknowledgments

This work was supported by donations from Haas Praezisionstechnik GmbH (Germany).

## Author Contributions

N.F. und S.B. initiated the project and prepared the manuscript. N.F. conceived and designed experiments. T.O. generated rat lines. N.F., F.K. and M.M. performed experiments. N.F. and M.M. analysed data. F.K. contributed NPCs and iPSC-derived neurons. T.O. and T.G. contributed reagents/ materials/ analysis tools.

## Supplementary data

**Supplementary fig. 1. Animal models**. (a) Position of the genomic deletion in LRRK2 k.o. rat. (b) Expression of human LRRK2 p.G2019S-transgene in different tissues of hG2019S tg rats. Whole brain lysates from a wild-type rat are used as negative control. The hLRRK2-specific signal is outlined in red.

**Supplementary fig. 2. Expression and phosphorylation of Akt in NPCs**. (a) Western Blot analysis of Akt protein expression and phosphorylation on T308 in L1-1 (left) und L2-2 NPCs (right). (b) Quantification of previous Western blot data for L1-1 and (c) L2-2 NPCs (n = 2 independent experiments).

**Supplementary table 1. Clinical data of LRRK2-associated PD patients**.

