## Supplemental data for "The Parkinson’s disease associated Leucine-rich repeat kinase 2 affects expression of Transferrin receptor 1 and phosphorylation of key signaling proteins in human iPSC-derived dopaminergic neurons"

Supplementary Figure 1.

A

exon 2 | intron II

...AAAAATGTCCACGTGCCTCTGTTGATAGTCCTGGACTCCTAC**ATGAGAG**TCGCCAGTGTGCAGCAGctaaaccgg...

7 bp deletion

B

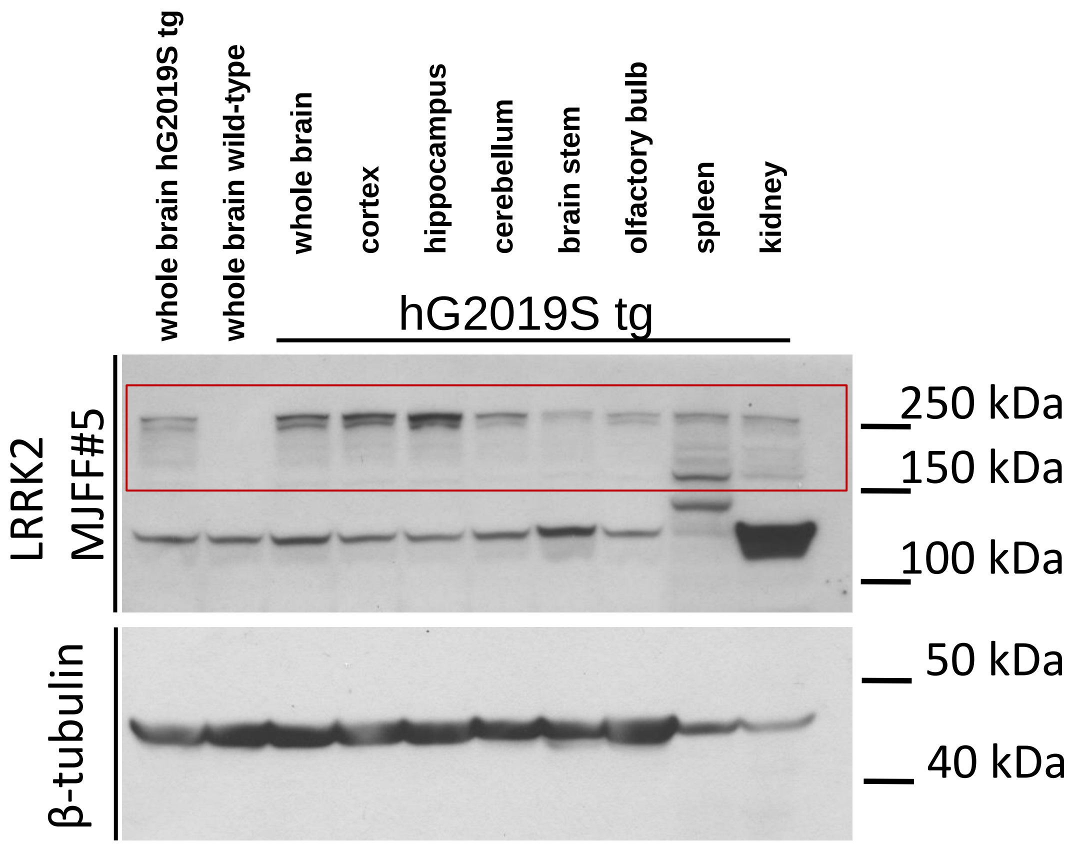

Supplementary Figure 2.

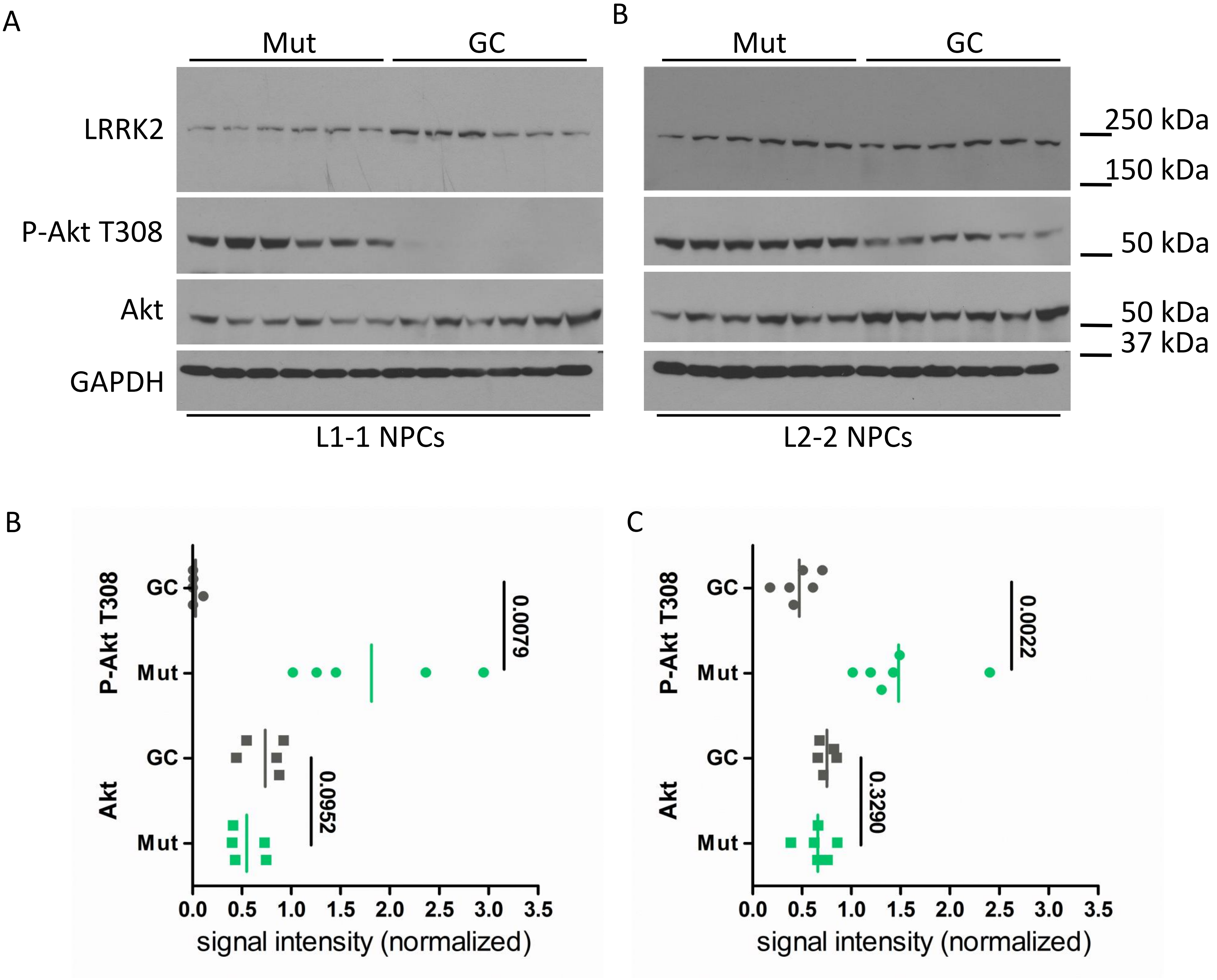

Supplementary Table 1.

| iPSC ID | genetic variant | age of onset | age at study | disease duration | sex | UPDRS III |
|---|---|---|---|---|---|---|
| L1 | *LRRK2* c.6055G>A (G2019S) | 40.36 | 50.95 | 10.59 | f | 13 |
| L2 | *LRRK2* c.6055G>A (G2019S) | 70.55 | 80.14 | 9.59 | f | 44 |

Clinical data of LRRK2-associated PD patients
